# Differential Effects of Haloperidol on Neural Oscillations During Wakefulness and Sleep

**DOI:** 10.1101/2024.06.17.599401

**Authors:** Diego Gallo, Matias Cavelli, Santiago Castro-Zaballa, Juan Pedro Castro-Nin, Claudia Pascovich, Pablo Torterolo, Joaquín González

**Affiliations:** Unidad Académica de Fisiología, Facultad de Medicina, Universidad de la República, Montevideo, 11800, Uruguay; Department of Psychiatry, University of Wisconsin-Madison, Madison, WI 53719, USA; Department of Psychology, King’s College, University of Cambridge, Cambridge, CB2 3EB, United Kingdom; Brain Institute, Federal University of Rio Grande do Norte, Natal, RN 59056, Brazil

**Keywords:** 1/f, Dopamine, Aperiodic Activity, Power Spectra, Neural Oscillations, Sleep, Gamma, Alpha

## Abstract

The electrical activity of the brain, characterized by its frequency components, reflects a complex interplay between periodic (oscillatory) and aperiodic components. These components are associated with various neurophysiological processes, such as the excitation-inhibition balance (aperiodic activity) or interregional communication (oscillatory activity). However, we do not fully understand whether these components are truly independent or if different neuromodulators modulate them in different ways. The dopaminergic system has a critical role for cognition and motivation, being a potential modulator of these power spectrum components. To improve our understanding of these questions, we investigated the differential effects of this system on these components using electrocorticogram recordings in cats, which show clear oscillations and aperiodic 1/f activity. Specifically, we focused on the effects of haloperidol (a D2 receptor antagonist) on oscillatory and aperiodic dynamics during wakefulness and sleep. By parameterizing the power spectrum into these two components, our findings reveal a state-dependent modulation of oscillatory activity by the D2 receptor across the brain. Surprisingly, aperiodic activity was not significantly affected and exhibited inconsistent changes across the brain. This suggests a nuanced interplay between neuromodulation and the distinct components of brain oscillations, providing insights into the selective regulation of oscillatory dynamics in awake states.

## Introduction

The electrical activity of the brain is typically studied by decomposing the signal into its frequency representation and grouping different frequencies into predetermined bands of interest (Buzsáki and Draguhn, 2004). The power spectrum has been hypothesized to reflect a sum of both periodic (clear peaks in the spectra) and aperiodic components (1/f power spectra decay; Freeman, 2000; Voytek and Knight, 2015; Donoghue et al., 2020, 2022). The aperiodic component likely arises from background neural activity and has been related to the excitation-inhibition balance (Gao et al., 2017), scale-free dynamics (He, 2014), and criticality (Bak et al., 1988). On the other hand, the periodic (oscillatory) component is driven by oscillatory activity at specific frequencies; for instance, gamma or theta oscillations (Buzsáki, 2002; Tort et al., 2009; Buzsáki and Wang, 2012; Gonzalez et al., 2023). This component is associated with the formation of cell-assemblies (Harris et al., 2003; Buzsáki, 2010) and interregional communication (Fries, 2005, 2015).

The grouping of frequencies into bands poses several problems. First, the limits of the studied bands are often chosen based on historical reasons without considering the mechanisms that generate a given oscillation (Lopes-Dos-Santos et al., 2018; Fernandez-Ruiz et al., 2023). Under this scenario, problems can arise since an oscillatory circuit may change frequencies, for instance, during different behaviors, animal models or experimental conditions (Tort et al., 2018; Kropff et al., 2021). Moreover, aperiodic parameters (exponent and offset) influence the power spectrum across all frequency bands (Voytek et al., 2015; Donoghue et al., 2020, 2022). Thus, changes in these parameters (e.g., an exponent increase) may affect different bands without truly reflecting changes at the oscillatory circuit level. Therefore, these confounding variables should be considered when analyzing the power spectrum and its changes during experimental manipulations.

An open question in the field is whether the periodic and aperiodic components of the power spectrum are differentially regulated, for instance, through neuromodulation. Understanding the neurochemical modulation of EEG power spectrum components is crucial for elucidating the underlying mechanisms of various neurological disorders and pharmacological interventions. The dopaminergic neurons of the ventral tegmental area (VTA) and substantia nigra pars compacta (SNpc) are considered a component of the activating system, and have a critical role in cognition and motivation (Nieoullon and Coquerel, 2003; Wise, 2004; Torterolo, 2019). Previous studies employing neurotoxic lesions of the SNpc (Parkinson’s disease model) lead to changes in the oscillatory component, increasing beta power and decreasing high-gamma activity (Cavelli et al., 2019; Kim et al., 2022). Furthermore, these conditions also affect the aperiodic component as well (Kim et al., 2022; Wang et al., 2022; Zhang et al., 2022; Wiest et al., 2023), reducing both the offset and the exponent (Kim et al., 2022). Interestingly, these effects on the aperiodic components were partially reversible with levodopa administration (Kim et al., 2022).

In this work, we recorded the electrocorticogram (ECoG) of five cats that exhibit clear oscillations and aperiodic 1/f activity (Cavelli et al., 2020). We then parametrized their power spectrum into the periodic and aperiodic components and studied the selective effects of haloperidol, a D2 dopamine receptor antagonist. Our results show that haloperidol differentially affects oscillatory dynamics across the brain in a state-dependent manner. Furthermore, while the specific aperiodic parameters (exponent and offset) were mildly affected by haloperidol, these changes compensated and caused aperiodic activity to remain constant. These results, based on the dopaminergic system, suggest that different neuromodulators can selectively regulate the periodic or the aperiodic oscillatory brain dynamics.

## Results

### Parametrizing the cat power spectrum into its periodic and aperiodic components

We recorded the ECoG of five awake adult cats in head-fixed conditions (Fig. 1A). Raw and filtered recordings already display noticeable alpha and gamma oscillations on a background of irregular activity (Fig. 1B). Since the power spectrum exhibits a mixture of aperiodic components, arising from background neural activity, and periodic components, reflecting oscillatory activity at specific frequencies, we sought to extract and analyze both components individually. For this purpose, we parameterized the raw power spectrum through the FOOOF (Fitting Oscillations and One-Over F) toolbox to separate and model both components (Donoghue et al., 2020). Briefly, the raw power spectrum (left panel in Fig. 1C) is initially fitted to estimate the aperiodic component, which is then subtracted from the raw spectrum. Peaks are identified based on the residual spectrum (aperiodic subtracted) employing a threshold, then a Gaussian is fitted around each peak. This process repeats until all points fall below the threshold. A multi-Gaussian fitting is performed on the residual spectrum, with the number of peaks determined in the previous step. The multi-Gaussian model is subtracted from the raw power spectrum, and a new aperiodic fit is computed. Next, the re-fitted aperiodic component (dashed line in the middle panel of Fig. 1C) is combined with the multi-Gaussian model to yield the final model of the power spectrum (solid line in the middle panel of Fig. 1C). Finally, we employed the multi-Gaussian model to study the periodic component exclusively (i.e., the oscillatory component, shown in the right panel in Fig. 1C). Notably, the parameterized model robustly captures the variance of the raw power spectrum (R^2^ = 0.99), indicating that 99% of the original power spectrum is reproduced by the model. Note that similar successful parameterizations are achieved in all five cats (Fig. S1).

**Figure 1.**
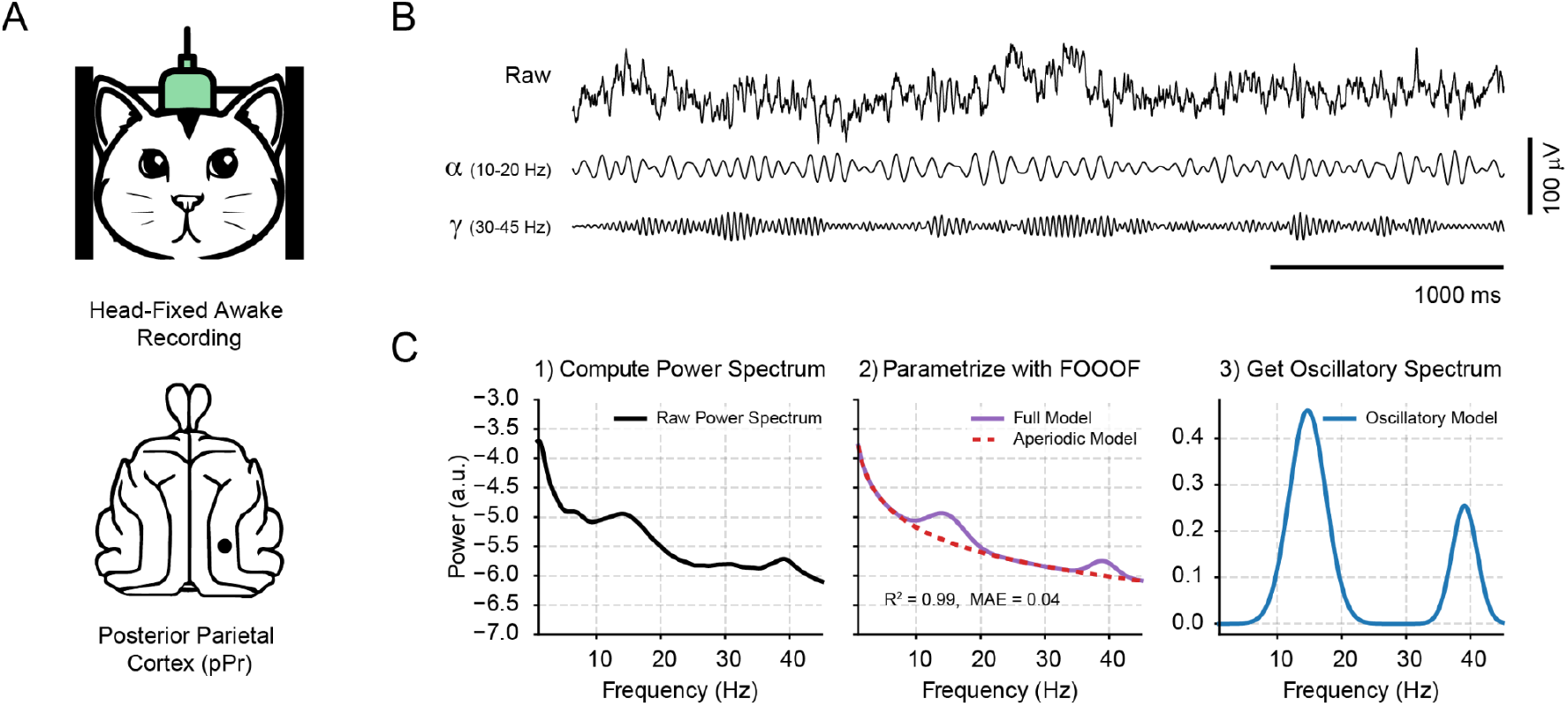
Decomposing the power spectrum into its periodic and aperiodic components. **A** Schematic representation of a cat recorded in awake, head-fixed conditions and the location of the right parietal posterior cortex (pPr). **B** Raw and filtered recordings (alpha: 10-20 Hz, gamma: 30-45 Hz) from the pPr of a representative animal during wakefulness. **C** Left: Representative power spectrum of the pPr cortex. All awake 10 seconds epochs within the first two hours of the recording session were employed. Middle: Parametrization of the power spectrum through the FOOOF framework. The aperiodic model is shown as a dashed line, while the full model (aperiodic + periodic components) is depicted by a solid line. Note that the model robustly captures the variance of the power spectrum (R^2^ = 0.99, i.e., 99% of the real power spectrum is reproduced by the FOOOF model). MAE: mean absolute error. Right: Periodic component extracted from the raw power spectrum. It’s worth mentioning that this component can be interpreted as the excess power above the aperiodic component.

### Haloperidol selectively promotes oscillatory activity

To study the effects of dopaminergic regulation of cortical dynamics, we administered haloperidol (4 mg/kg i.m.) in awake head-fixed cats and recorded the first two hours after haloperidol achieved maximal concentration in the brain (Fig. 2A, Froemming et al., 1989). We then fitted the resulting power spectrum with the FOOOF model and achieved a satisfactory parametrization judged by the goodness of fit metrics for one cortical area shared across all cats (posterior Parietal Cortex, pPr) (R^2^ > 0.9 and mean absolute error (MAE) < 0.1 for all animals, Fig. 2B). Note there were no significant differences either in the R^2^ or MAE between haloperidol and the basal condition (no drug administered; R^2^: t(4) = 1.20, p = 0.29; MAE: t(4) = 1.16, p = 0.31).

**Figure 2.**
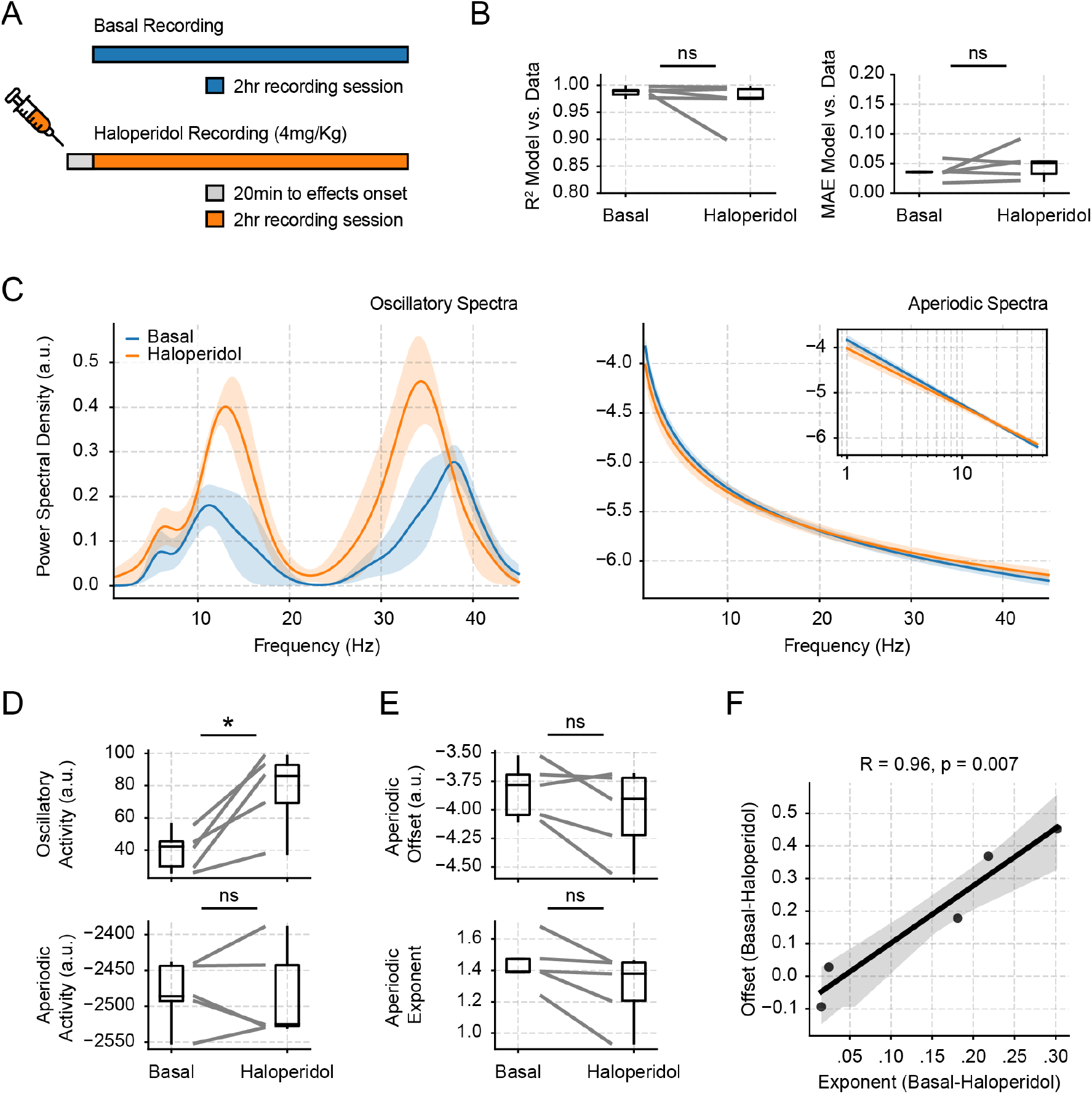
Haloperidol selectively affects oscillatory activity during wakefulness. **A** Schematic representation of the experimental protocol. **B** Goodness of fit metrics (R^2^ and MAE) for the FOOOF parameterization in all animals during both recording conditions for the pPr cortex; ns, non-significant. **C** Left: Mean oscillatory power spectrum (± SEM; n = 5 cats) during basal and haloperidol recordings. Right: Mean aperiodic power spectrum (± SEM; n = 5 cats) during basal and haloperidol recordings. The inset displays a log-log representation of the same plot and visualizes the aperiodic component as a linear trend. **D** Activity of the periodic (top) and aperiodic (bottom) power spectrum components for both recording conditions. **E** Offset (top) and exponent (bottom) of the aperiodic model for both recording conditions. **F** Correlation between the offset and exponent differences (Basal-Haloperidol). Note that these parameters follow a linear relationship, in which the offset difference is compensated by the exponent changes, explaining why the aperiodic activity does not change between conditions.

Haloperidol selectively increases oscillations without changing the overall aperiodic activity. We decomposed the power spectrum into both components and quantified total oscillatory and aperiodic power by summing the power in the periodic/aperiodic component across all frequencies (referred to hereafter as oscillatory and aperiodic activity). Thus, an index of the activity of each component is obtained to compare across conditions. We found that haloperidol significantly increased oscillatory activity compared to the basal condition (Fig. 2C, D top; t(4) = 4.17, p = 0.01). In contrast, the aperiodic activity remained similar between haloperidol and the basal condition (Fig. 2C, D bottom; t(4) = 0.01, p = 0.99).

To gain further insights into why the aperiodic component did not change, we measured the two parameters that determine its decay: the aperiodic exponent (dictating the decay rate) and its offset (y-axis intercept). We found that haloperidol shows a tendency to decrease the exponent and offset (Fig. 2E) yet, these results were not significant (Exponent: t(4) = 2.65, p = 0.05; Offset: t(4) = 1.83, p = 0.14). Nevertheless, the changes in both parameters were significantly correlated (Fig. 2F; R(3) = 0.96, p = 0.007). This result explains why the overall aperiodic activity remained constant. Both exponent and offset changes compensated; i.e., when the exponent decreased, the offset also decreased, maintaining the aperiodic activity sum equal.

### Haloperidol oscillatory effects occur across cortical regions and the thalamus

To gain further insights into haloperidol effects on oscillatory dynamics, we compared its effects across all areas recorded during wakefulness. Each cat contained a different array of electrodes, spanning several neocortical sites in both hemispheres (prefrontal, motor, somatosensory, auditory, parietal, and visual areas), the olfactory bulb, and the lateral geniculate nucleus (Fig. 3A). Of note, the power spectrum parametrization was successful in all locations (R^2^ > 0.9 and MAE < 0.1 for all sites and all animals, Fig.S2).

**Figure 3.**
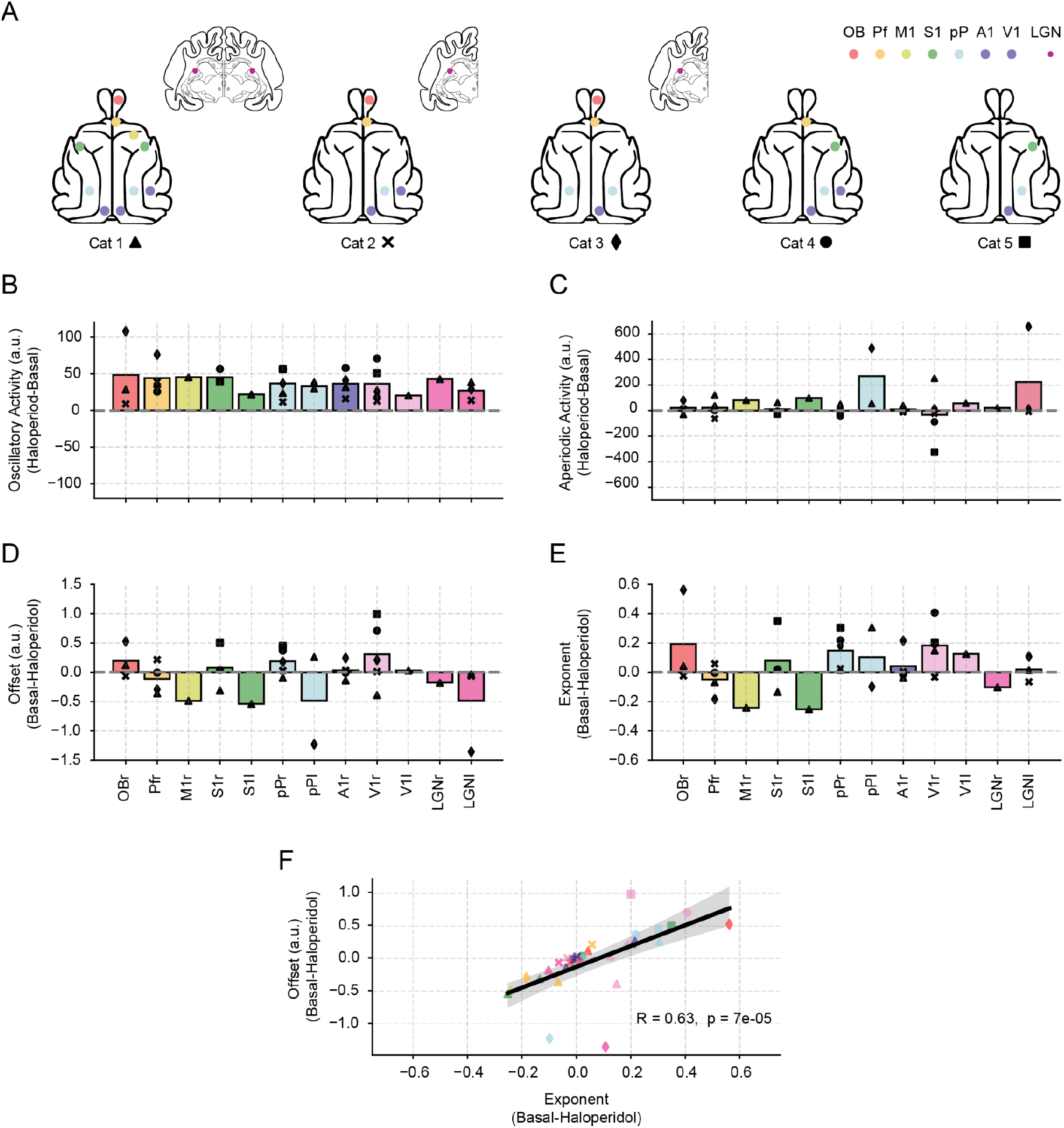
Brain-wide oscillatory changes induced by haloperidol during wakefulness. **A** Diagram of a cat’s brain, showing the different regions recorded in each of the five cats. Each area is color-coded, and each marker shows a different cat. **B** Bar graph representing the activity difference between haloperidol and basal conditions for the oscillatory activity. Each colored bar shows a region in the brain diagrams depicted in A. Individual points within each bar show an individual animal with a distinct symbol. The height of the bars shows the mean activity difference. **C** Same as in panel B but for the aperiodic component. **D** Offset changes induced by haloperidol for all recording sites. **E** Exponent changes induced by haloperidol for all recording sites. **F** Correlation between the offset and exponent differences (Basal-Haloperidol) for all electrodes and animals. OB: Olfactory Bulb; Pf. Prefrontal Cortex; M1: Primary Motor Cortex; S1:Primary Somatosensory Cortex; pP: posterior Parietal Cortex; A1: Primary Auditory Cortex; V1: Primary Visual Cortex; LGN: Lateral Geniculate Nucleus; r: right; l: left.

Haloperidol promotes oscillations across the brain. When comparing across all recording sites, we found that haloperidol significantly increased the periodic activity in all electrodes (Fig. 3B; Drug: F(1) = 35.68, p = 4.3 x 10^-7^; Area: F(11) = 4.62, p = 0.00013; Drug x Area: F(11) = 0.11, p = 0.99). On the other hand, the effects of haloperidol on the aperiodic activity were not significant across brain areas (Fig. 3C; Drug: F(1) = 1.17, p = 0.28; Area: F(11) = 5.83, p = 0.00001; Drug x Area: F(11) = 0.39, p = 0.95). This observation contrasts with the homogenous increases in periodic activity elicited by haloperidol.

To delve deeper into the lack of consistency for haloperidol effects on aperiodic activity, we analyzed the exponent and offset variation across the brain. Importantly, we found that the offset exhibited inconsistent changes across sites, increasing in some areas and decreasing in others (Fig. 3D; Drug: F(1) = 0.02, p = 0.88; Area: F(11) = 5.98, p = 9 x 10^-06^; Drug x Area: F(11) = 0.41, p = 0.94). The same occurred with the exponent (Fig. 3E; Drug: F(1) = 1.31, p = 0.26; Area: F(11) = 2.38, p = 0.02; Drug x Area: F(11) = 0.35, p = 0.96). Moreover, we also found that the changes in exponent and offset were significantly correlated for all recording sites (Fig. 3F; R(3) = 0.63, p = 7 x 10^-05^), evidencing the compensation of both parameters in the aperiodic component. Thus, these results suggest that haloperidol selectively affects periodic activity, since whole aperiodic activity did not change. Furthermore, while aperiodic parameters may change in some areas, these variations are inconsistent across the brain.

### Haloperidol oscillatory effects are absent during sleep

In order to explore whether the effects of haloperidol are state-dependent, we also studied its effects during natural non-NREM (NREM) sleep (Fig.4A). Note that we achieved a satisfactory parametrization of the pPr across all cats during sleep (R^2^ > 0.9, Fig.S3). First, we found that for the cortical location shared in all cats (pPr), haloperidol did not produce any significant differences in either the periodic or aperiodic components during this state (Fig. 4B, C; Oscillatory Activity: t(4) = 1.15, p = 0.31; Aperiodic Activity: t(4) = 1.96, p = 0.12). Note that there were no significant changes in either aperiodic offset or exponent (Fig. 4C; Offset: t(4) = 2.59, p = 0.06; Exponent: t(4) = 2.49, p = 0.07), even though we observed a decreasing tendency of these parameters during sleep.

**Figure 4.**
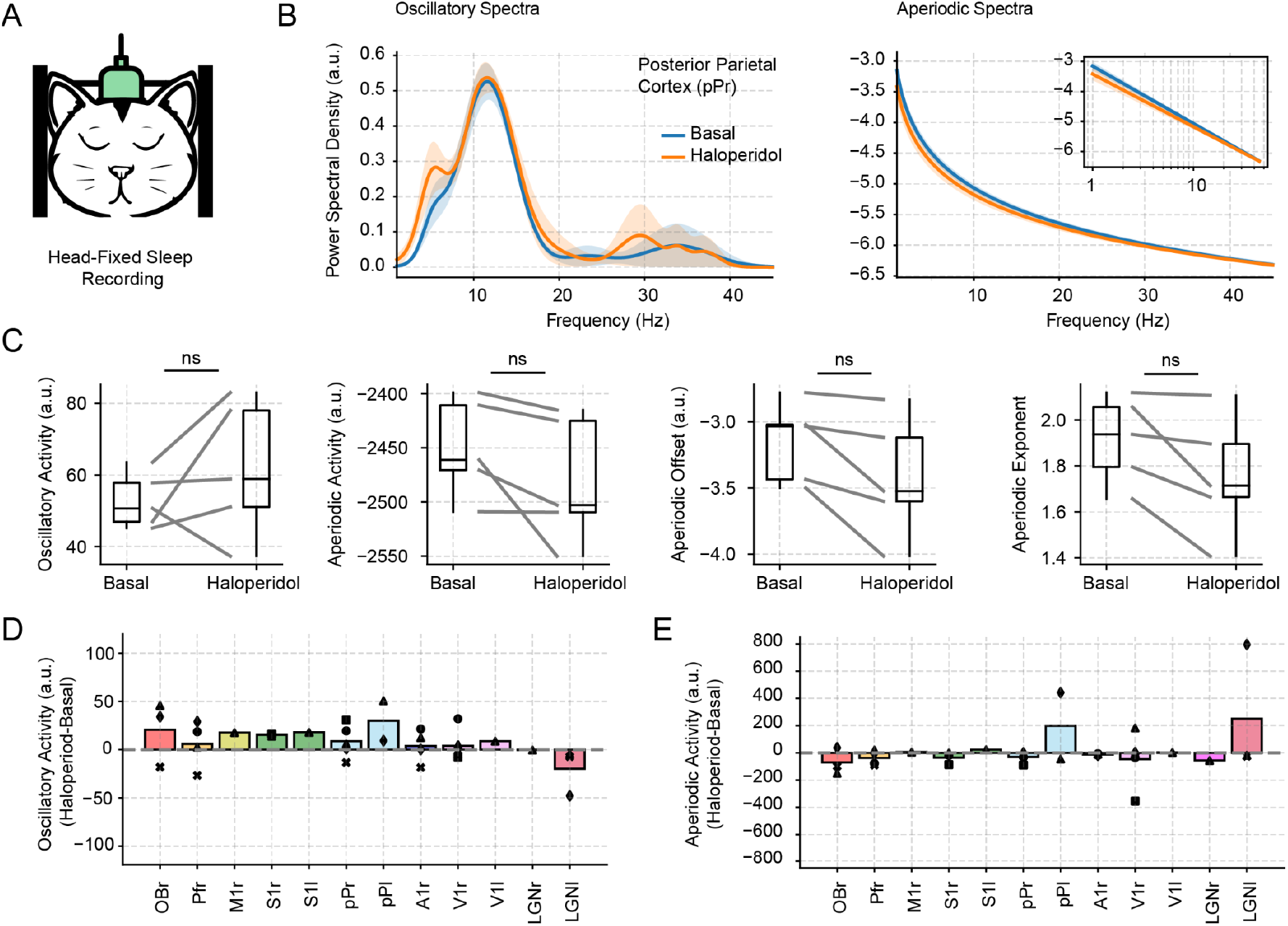
Haloperidol effects on oscillatory activity are absent during NREM sleep. **A** Illustration of a cat recorded during sleep in head-fixed conditions. **B** Left: Mean oscillatory spectra for the basal and haloperidol conditions (± SEM; n = 5 cats). Right: Mean aperiodic spectra for both conditions. **C** Periodic and aperiodic activity and aperiodic parameters (Offset and Exponent) for both recording conditions. ns, non-significant. **D** Bar graph representing the difference between haloperidol and basal conditions for the oscillatory activity. Each colored bar shows a region in the brain diagrams. Individual points within each bar show an individual animal with a distinct symbol. The height of the bars shows the mean difference. **E** Same as in panel **D** but for the aperiodic component.

Moreover, we also assessed these effects throughout the brain where we confirmed the lack of results almost across all recording sites for both the periodic (Fig. 4D; Drug: F(1) = 3.38, p = 0.07; Area: F(11) = 1.01, p = 0.46; Drug x Area: F(1) = 0.67, p = 0.76) and aperiodic components of the power spectrum (Fig. 4E; Drug: F(1) = 0.02, p = 0.88; Area: F(11) = 5.27, p = 3.7 x 10^-05^; Drug x Area: F(11) = 0.4, p = 0.94). Thus, these results suggest that haloperidol promotion of oscillatory activity depends on the behavioral state.

### Haloperidol enhances alpha and gamma activity during wakefulness

Finally, we sought to understand which specific oscillations were responsible for the periodic changes elicited by haloperidol during wakefulness. To this end, we employed a data-driven approach to define the narrow frequency bands of interest. Specifically, we collected all the power spectrum peaks for all animals and electrodes (suggestive of a true oscillation, Yuval-Greenberg et al., 2008) and measured their distribution. We observed three main frequency clusters for both basal and haloperidol conditions (Fig.5A left). These clusters corresponded roughly to the theta (4-9 Hz), alpha (10-20 Hz), and gamma (25-45 Hz) bands (note that this cluster also encompasses frequencies typically associated with the beta band). Haloperidol did not change the width of each narrow band oscillation (Fig.5A right) or the total number of peaks detected for the total range of frequency and for each frequency band (Fig.5 C; Theta: t(4) = 0.28; Alpha: t(4) = 0.17; Gamma: t(4) = 0.46; Total Peaks: t(4) = 0.19). Note the number of peaks detected in the theta band was inconsistent across animals (and also across cortical recordings sites), thus we will not consider this band further in the analysis.

To study the narrow-band power of each of these three bands, we chose to compare the peak power of alpha and gamma oscillations between basal and haloperidol conditions (Fig. 5D), thus mitigating the effects of possible frequency shifts. Nonetheless, our results also hold when comparing the power in the whole band (Fig.S4). Notably, we found that haloperidol significantly increased alpha and gamma power compared to the basal recording (Fig. 5D; Alpha Peak Amplitude: Drug: F(1) = 28.58, p = 2.3 x 10^-5^; Area: F(11) = 2.76, p = 0.02; Drug x Area: F(11) = 0.51, p = 0.88; Gamma Peak Amplitude: Drug: F(1) = 14.29, p = 0.0005; Area: F(11) = 4.46, p = 0.0002; Drug x Area: F(11) = 0.05, p = 0.99). When comparing the peak frequency in each band, we found that haloperidol did not significantly shift alpha rhythms (Fig. 5E; Alpha Shift Frequency: Drug: F(1) = 0.13, p = 0.72; Area: F(11) = 0.62, p = 0.79; Drug x Area: F(11) = 0.22, p = 0.99). In contrast, haloperidol significantly decreased the frequency of gamma oscillations (Fig. 5E; Gamma Shift Frequency: Drug: F(1) = 28.75, p = 3 x 10^-6^; Area: F(11) = 0.43, p = 0.94; Drug x Area: F(11) = 0.18, p = 0.99). Thus, our results show that haloperidol increased the power of the oscillations while it selectively decreases gamma frequency.

**Figure 5.**
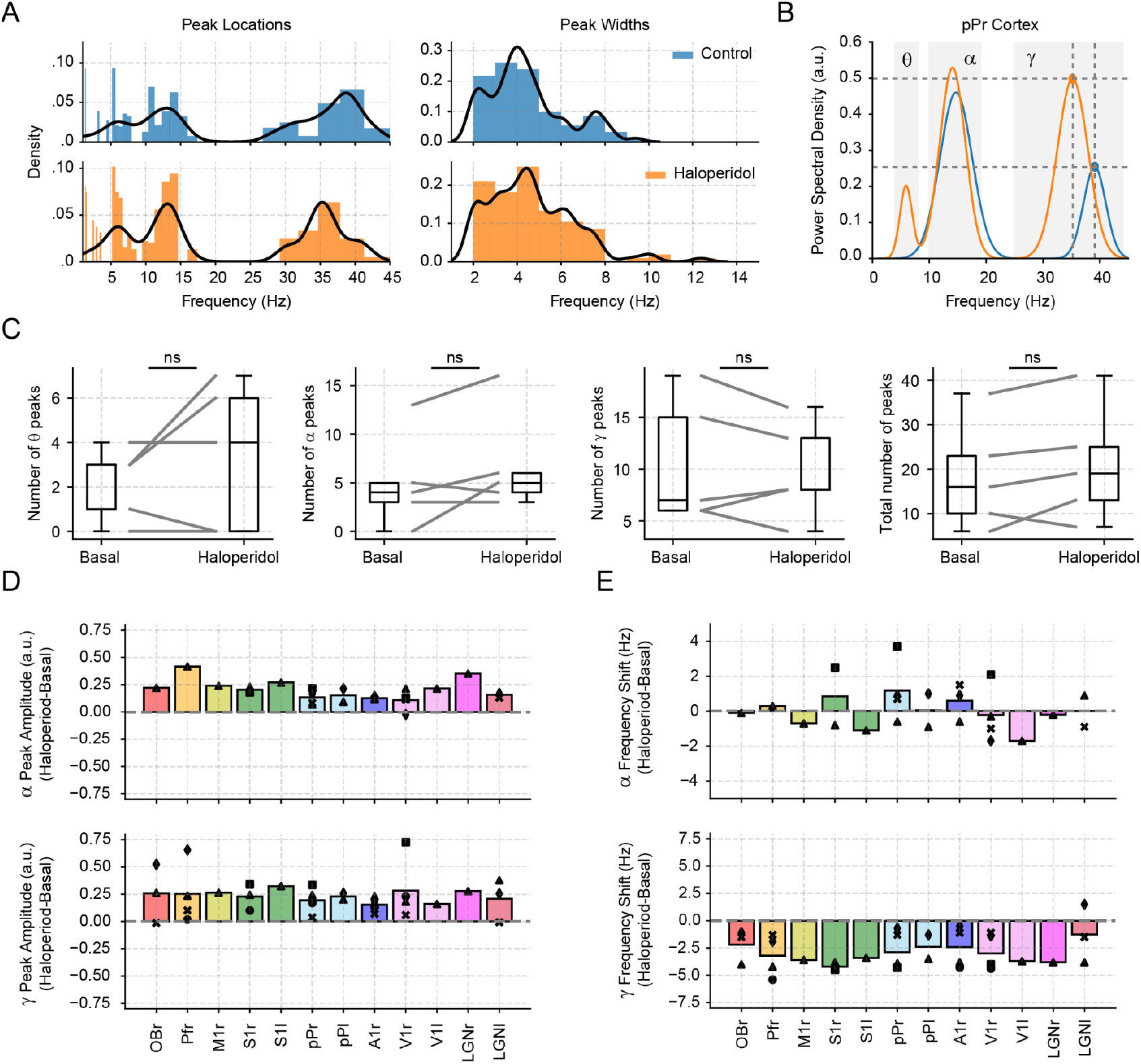
Band-limited effects of haloperidol administration during wakefulness. **A** Left: Distribution of peak frequency locations for all electrodes and animals during basal and haloperidol recordings. All peaks in the spectrum were detected through FOOOF. Note that three main clusters of frequencies appear, corresponding to the theta, alpha, and gamma frequency bands. We employed logarithmic frequency bins to construct the histogram; thus the width of each bin increased from the lower frequencies (higher bin resolution) to the higher frequencies. Right: Distribution of peak widths detected. **B** Periodic spectrum of the pPr cortex from a representative animal during both conditions. Dashed lines and dots depict the peak amplitude and frequency of the gamma frequency band. **C** Number of peaks detected through FOOOF for the whole power spectrum, and divided for each frequency band for basal and haloperidol recordings. **D** Bar graph representing the peak amplitude difference between haloperidol and basal conditions for the three frequency bands detected. Each colored bar shows a region in the brain diagrams. Individual points within each bar show an individual animal with a distinct symbol. The height of the bars shows the mean difference. **E** Same as in panel **D** but showing the peak frequency changes between both conditions.

## Discussion

Here, we showed that haloperidol selectively affects oscillatory dynamics across the brain in a state-dependent manner i.e., these oscillatory changes only occurred during wakefulness but not during NREM sleep. Specifically, haloperidol increased the power of alpha, and gamma bands, reducing the peak frequency of the latter. Notably, while haloperidol tended to affect the exponent and offset of the aperiodic component, these changes were not consistent across areas and even compensated, making the total aperiodic power remain constant.

A noticeable result we obtained was the differential effects of haloperidol on periodic and aperiodic power. Our results suggest that dopaminergic transmission through the D2 receptor may affect oscillatory activity exclusively without changing the brain’s aperiodic activity. Therefore, we deem it likely that both power spectrum components get regulated independently since their origins likely involve different neural substrates. Note that the circuits underlying oscillations are better known and tested, for instance excitatory-inhibitory loops in the neocortex (Wang, 2010; Buzsáki and Wang, 2012) or in the thalamus (Golomb et al., 1994; Wang et al., 1995). Conversely, theoretical models suggest that the universality class of the network (for instance, a branching model) might be the defining element that dictates the types of spiking patterns captured by the aperiodic component (Brake et al., 2024). In further support of our hypothesis, Wang et al., showed that both periodic and aperiodic parameters are more closely correlated with each other than across them.

Furthermore, since the extracellular levels of dopamine change with arousal and are minimal during NREM sleep (Léna et al., 2005), we considered it important to assess haloperidol effects during this state. Consistently, we found that the oscillatory changes previously observed during wakefulness disappeared during NREM sleep, suggesting that the total dopamine levels were reduced in this state and were linked to the regulation of periodic activity.

Several of our results are in close agreement with both preclinical models of Parkinson’s disease and clinical patients. We observed a non-significant trend of the exponent and offset in the pPr to decrease. Similar results occur in preclinical 6-OHDA models in the subthalamic nucleus (Kim et al., 2022); and also in Parkinson patients when comparing on vs. off levodopa administration (Wang et al., 2022; Zhang et al., 2022; Wiest et al., 2023). Interestingly, similar to our topographical results, (Wang et al., 2022) also reported heterogeneous offset and exponent differences between healthy individuals and Parkinson’s patients without treatment. In addition, we found that gamma oscillations decreased in frequency and shifted into the beta range (15-30 Hz), consistent with previously reported beta oscillation increases (Cavelli et al., 2019; Kim et al., 2022).

Moreover, our results contrast those obtained in schizophrenic patients, further supporting the antipsychotic properties of haloperidol. For instance, patients with schizophrenia showed an increase in the aperiodic exponent (Peterson et al., 2023), which is consistent with the haloperidol tendency to decrease this parameter. Schizophrenic patients also show a reduction of gamma activity (Oribe et al., 2019; Hirano and Uhlhaas, 2021), consistent with the gamma increase we observed.

Neuroscience has traditionally focused on oscillatory activity (Adrian, 1942; Buzsáki and Draguhn, 2004; Cole and Voytek, 2017; González et al., 2023), but we need to further understand how aperiodic activity arises and how to study it. Notably, the exponent and the offset exert opposite contributions to the overall signal. Reductions in the exponent (dictating the aperiodic decay) cause the spectrum to become flatter, thus increasing total power; while reductions in the offset (aperiodic power at f=0) decrease the overall power since the power function starts at a lower value. A surprising result we obtained was the lack of effects of haloperidol on the aperiodic power. We found that the variations in both parameters compensated, making the aperiodic component remain constant. This result agrees with previous works (Podvalny et al., 2015; Donoghue et al., 2020; Kim et al., 2022; Wang et al., 2022; Zhang et al., 2023).

Therefore, these results raise several questions about how to study aperiodic activity. 1) What parameters should be studied? 2) Should we only consider the exponent value since it has a more defined biological correlation (excitatory-inhibitory balance)? 3) What do the changes in the offset mean from a neural network perspective? For these reasons, we raise caution, since several studies have consistently reported aperiodic slope changes but have not leveraged the influence of the offset, raising the question of whether aperiodic activity actually changes or not (Salvatore et al., 2024). This lack of offset control may lead to potential misinterpretation of the results (that aperiodic activity changes when perhaps it does not). Thus, we argue that until we have a stronger theoretical and biological interpretation of these parameters, both of them should be reported.

Several limitations of our study should be acknowledged. First, we employed haloperidol to understand if dopaminergic neuromodulation selectivity regulates the periodic and aperiodic power spectrum components. Importantly, haloperidol is not a completely selective D2 antagonist and also interacts (with much lower affinity) with serotonergic, alpha-adrenergic, histaminergic, or cholinergic receptors (Ohta, 1976; Navailles et al., 2006; Beach et al., 2020). Second, we note that our study lacks a sham control; our control was a basal recording and not the administration of the vehicle. Yet, pilot experiments by our group showed that the systemic administration of saline solution does not influence brain dynamics, thus, we have consistently employed basal recordings as controls (Castro-Zaballa et al., 2018, Castro-Zaballa 2019). Third, given that we administered haloperidol systemically, we cannot assert if the differential periodic and aperiodic regulation occurs locally (intracortical regulation of the local circuitry) or globally (promoted by subcortical centers).

## Methods

### Animals

Five adult cats were included in this study, some of which were also utilized in prior research (González et al. 2023; Cavelli et al. 2020). The Institutional Animal Care Facility assessed these animals as being in good health. All experimental procedures adhered to the Guide for the Care and Use of Laboratory Animals (8th edition, National Academy Press, Washington DC, 2011) and received approval from the Institutional Animal Care Commission (No: 070153-000089-17). Measures were taken to minimize pain, discomfort, or stress, and effort was made to employ the minimum number of animals necessary to generate reliable scientific data.

### Surgical Procedure

The animals underwent surgical implantation of electrodes to monitor their sleep/wake states, as described in our previous work (Castro-Zaballa et al., 2018; Cavelli et al., 2020). Before anesthesia, each cat was administered premedication consisting of xylazine (2.2 mg/kg, i.m.), atropine (0.04 mg/kg, i.m.), and antibiotics (ceftriaxone 50 mg/kg, i.m.). Anesthesia was induced with ketamine (15 mg/kg, i.m.) and maintained with a gas mixture of isoflurane in oxygen (1%–3%). The animals’ heads were secured in a stereotaxic frame, and stainless-steel screw electrodes (1 mm diameter) were placed on the surface of various cortical regions. Bipolar electrodes were also implanted in the lateral geniculate nuclei (LGN). All electrodes were connected to a Winchester plug affixed to the skull with acrylic cement to ensure stable positioning without causing discomfort. Following surgery, the animals were allowed to recover and adapt to the recording environment for at least two weeks.

### Polysomnographic Recordings

Experimental sessions lasting 2 hours were conducted between 11 a.m. and 1 p.m. in a temperature-controlled environment (21–23°C). During these sessions, the animals’ heads were held in a stereotaxic position with four steel bars inserted into the chronically implanted plastic tubes as the body rested in a sleeping bag. The electrocorticogram (ECoG) was recorded with a monopolar configuration, utilizing the same reference electrode in the left frontal sinus. Bioelectric signals were amplified (×1000), filtered (0.1–500 Hz), sampled (1024 Hz, 16 bits), and stored in a PC using Spike 2 software (CED). Data were collected during spontaneously occurring wakefulness (W), non-REM (NREM) sleep, and REM sleep.

### Haloperidol Administration

We conducted recordings under basal conditions without pharmacological manipulation and 20 minutes after administering 4 mg/kg of haloperidol. This dosage selection was based on prior studies in the cat aimed at inducing maximal effects (Monti, 1968).

### Classification of sleep-wake states

The obtained recordings were classified into Wakefulness (W), NREM sleep, and Rapid Eye Movement (REM) sleep in epochs of 10 seconds. W was identified by the presence of active EEG, characterized by low amplitude and high frequency of electroencephalographic waves, accompanied by high amplitude in the EMG. NREM sleep was identified through an EEG characterized by high-amplitude slow waves (0.5-4 Hz) and sleep spindles (11-15 Hz), accompanied by a decrease in EMG amplitude. Lastly, REM sleep exhibits low-amplitude, high-frequency EEG similar to W, with the particularity of muscle atonia observed in the EMG. We did not analyze REM sleep because the number epochs in this state during the first 2-hours of recording was small.

### Spectral analysis of the electrographic signals

For all analyses, we used Python 3 with numpy (https://numpy.org/), scipy (https://docs.scipy.org/), matplotlib (https://matplotlib.org/), FOOOF (https://fooof-tools.github.io/fooof/).

We obtained spectral power using the welch function in scipy (parameters: window = 1024, noverlap = [], fs = 1024, nfft = 10240), corresponding to a 1-second window, with a 0.5-second overlap and a frequency resolution of 0.1 Hz. Since the power spectrum exhibits a mixture of aperiodic components, representing background neural activity, and periodic components that indicate oscillatory activity at specific frequencies, we sought to extract and analyze both types of components individually.

To achieve this, we employed the FOOOF Python toolbox. FOOOF (Fitting Oscillations and One-Over F) is a technique that allows us to parameterize power spectra and extract their periodic and aperiodic components. First, the raw power spectrum decay is fitted to estimate the aperiodic component. Second, the aperiodic component gets subtracted from the raw spectrum, yielding a residual mixture of periodic oscillatory peaks and noise. Third, peaks are found based on a threshold (2 standard deviations in our case), and a Gaussian is fitted surrounding this peak. This fitted Gaussian is subtracted from the residual spectrum. This process is repeated until all points fall below a noise threshold. Fourth, a multi-Gaussian fitting is then performed on the residual power spectrum employing the number of peaks found in the previous step, i.e., if four peaks are detected, a four-peak multi-Gaussian distribution is fitted. Fifth, the multi-Gaussian model is subtracted from the raw power spectrum, and a new aperiodic fit is computed. Sixth, the re-fitted aperiodic component is combined with the multi-Gaussian model to give the final model of the power spectrum. Finally, we evaluated the goodness of fit by computing 1) the coefficient of determination (R^2^) and 2) the mean absolute error (MAE) between the FOOOF model and the raw power spectrum.

For this work, we considered the power spectrum exclusively between 1 and 45 Hz. We made the fitting using the ‘fixed’ aperiodic mode because there was no distinct ‘knee’ in the power spectrum upon visual inspection in the logarithmic scale (i.e., the signal exhibited a nearly linear pattern across the designated frequency range). Additionally, we set the peak width limits to [2, 20] while leaving the values of max_n_peaks and min_peak_height at their default settings.

To compare the periodic and aperiodic power spectrums, we summed the power in each component across all frequencies to obtain a single value. These values are referred to as periodic and aperiodic activity in the figures and text.

### Statistics

We show group data as either mean ± SEM or regular boxplots showing the median, 1st, and 3rd quartiles, and the distribution range without outliers. We employed paired two-tailed t-test to compare between basal and haloperidol, and Two-Way ANOVA to compare the effects of haloperidol across all electrodes. We set p<0.05 to be considered significant.

## Author contributions

D.G., S.C.Z., P.T., and J.G. designed the experiments; D.G., M.C., S.C.Z., J.P.C. and C.P. conducted the experiments; D.G. and J.G., wrote analysis software and analyzed the data; D.G., M.C., S.C.Z., J.P.C., C.P., P.T., and J.G. were involved in the discussion and interpretation of the results; D.G., P.T., and J.G. wrote the manuscript. All the authors participated in the critical revision of the manuscript, added important intellectual content, and approved the final version.

## Acknowledgement

This study was supported by the “Programa de Desarrollo de Ciencias Básicas, PEDECIBA” (https://www.pedeciba.edu.uy/es/area/biologia/), Agencia Nacional de Investigación e Innovación (ANII), and the “Comisión Sectorial de Investigación Científica” (CSIC) I + D grupos 2022-22620220100148 grant from Uruguay (https://www.csic.edu.uy/). The funders had no role in study design, data collection and analysis, decision to publish, or preparation of the manuscript.

## Supplementary Material

**Figure S1.**
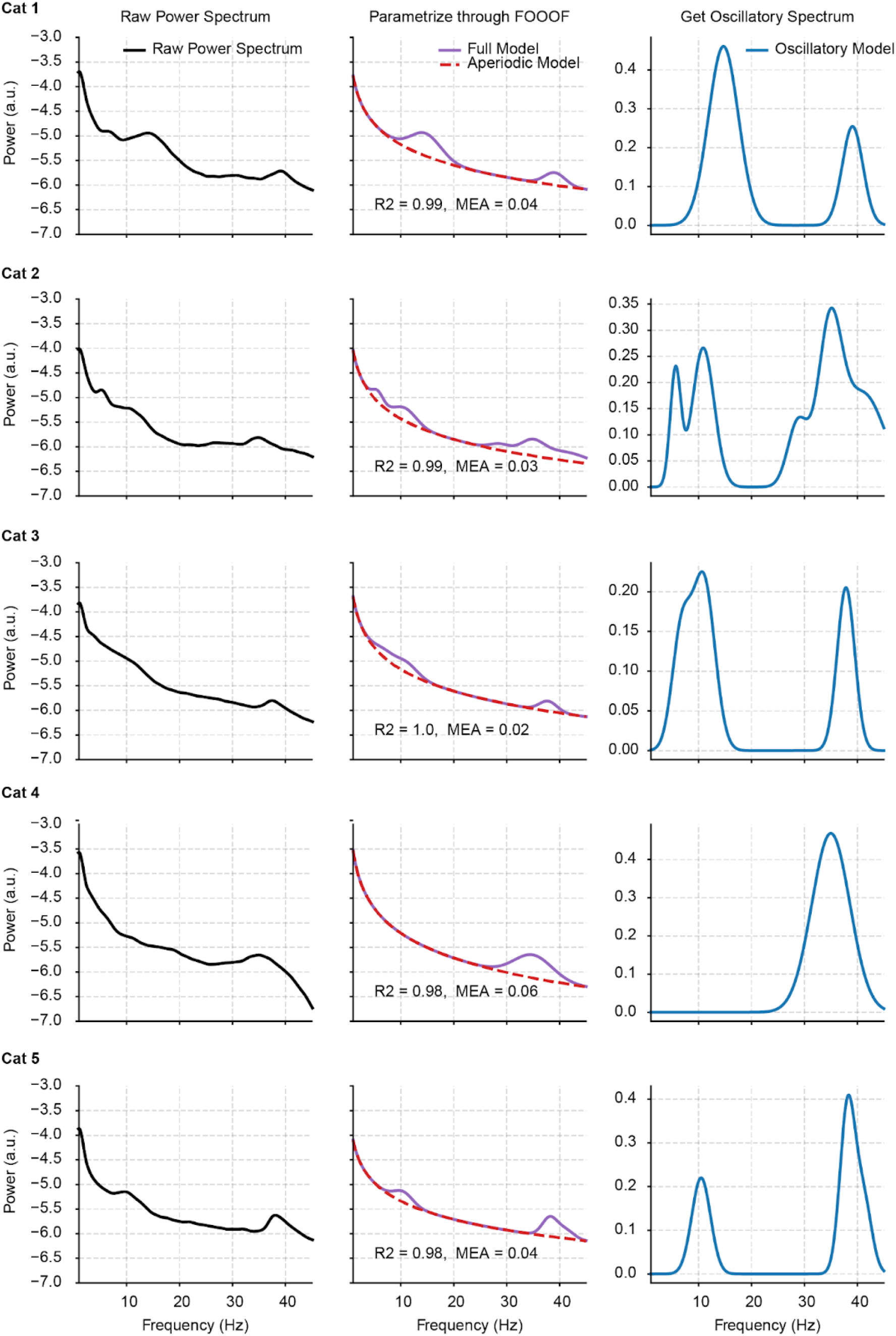
Parametrizing the power spectrum in all animals. Left: Representative power spectrum of the pPr cortex in basal conditions during wakefulness. All awake epochs within the first two-hours of the recording session were employed. Middle: Parametrization of the power spectrum through the FOOOF toolbox. The aperiodic model is shown as a dashed line while the full model (aperiodic + periodic components) is depicted by the solid line. R^2^: coefficient of determination. MAE: mean absolute error. Right: Periodic component extracted from the raw power spectrum. It’s worth mentioning that this component can be interpreted as the excess power above the aperiodic component. The differences among the animals in the periodic component are probably due to their level of alertness. Note that gamma activity in the cat is highly dependent of this cognitive function (Castro et al., 2013), while alfa power increases in quiet wakefulness (Rougeul-Buser and Buser, 1997; Crochet and Sakai, 1999).

**Figure S2.**
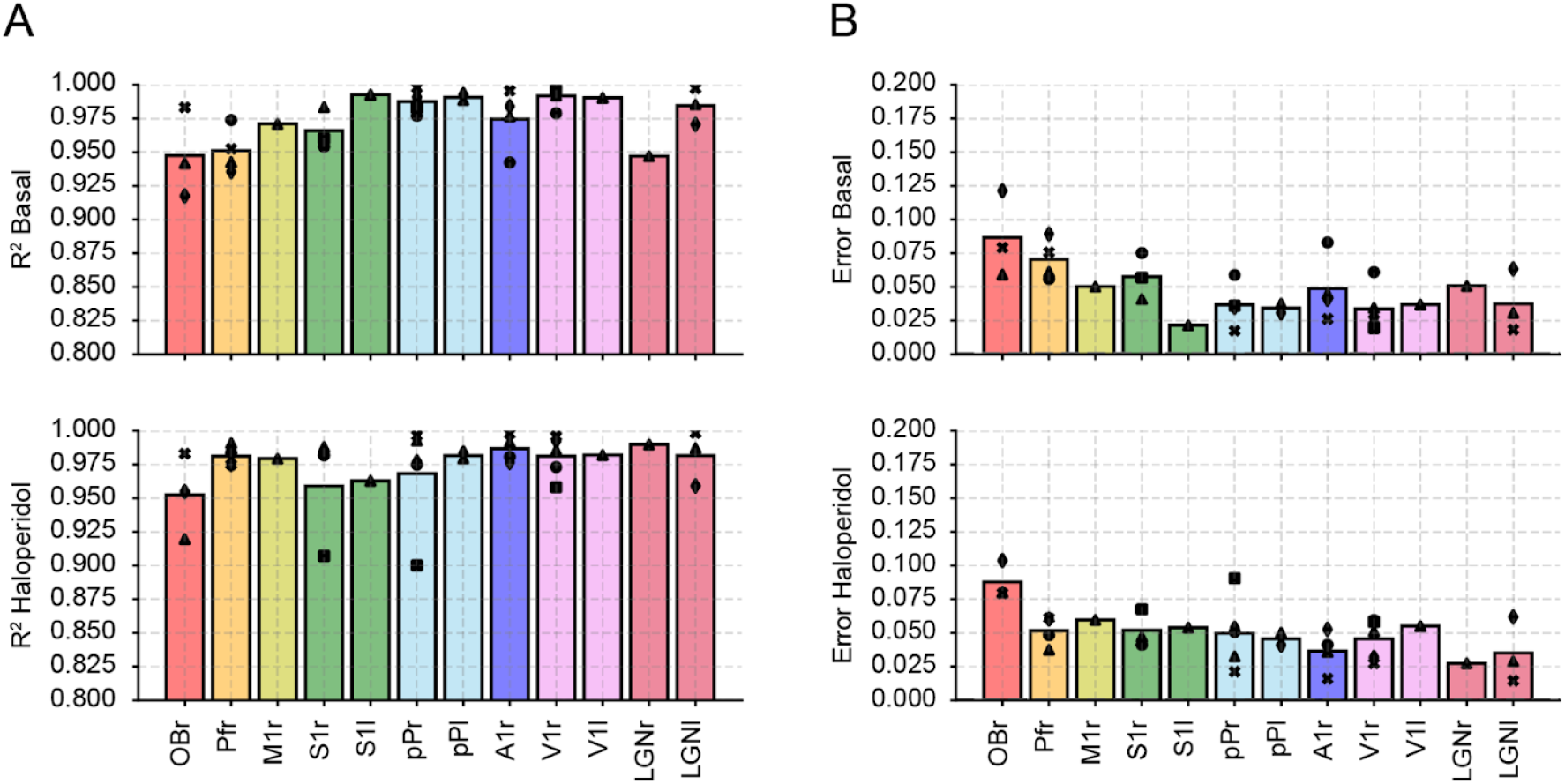
Assessing FOOOF model fit in all recording sites. **A-B** Goodness of fit metrics (R^2^ and MAE) for the FOOOF parameterization in all animals and all recording sites during basal and pharmacological conditions during wakefulness. Each colored bar shows a region in the brain diagrams represented in the figure 3A. Individual points within each bar show an individual animal with a distinct symbol, also shown in figure 3A.

**Figure S3.**
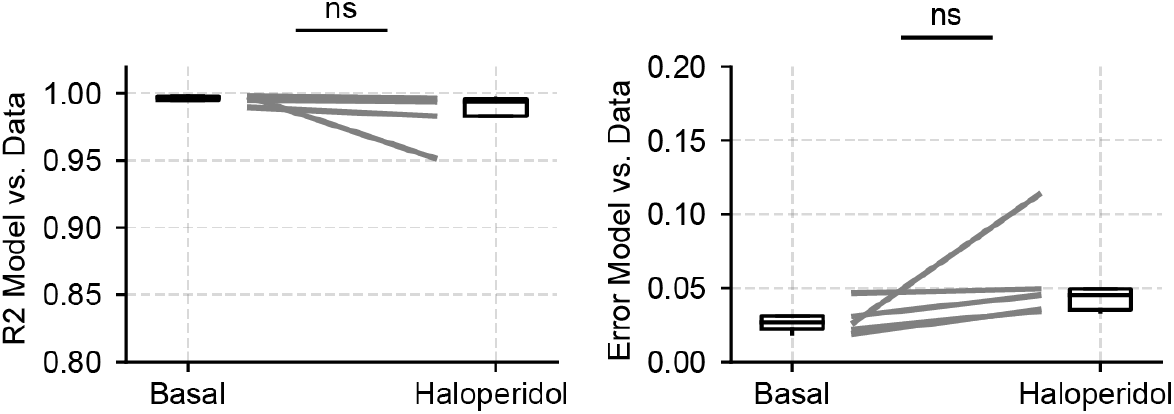
Assessing FOOOF model fit during NREM. Goodness of fit metrics (R^2^ and MAE) for the FOOOF parameterization for sleep recordings in all animals during both basal and pharmacological conditions for the pPr; ns, non-significant.

**Figure S4.**
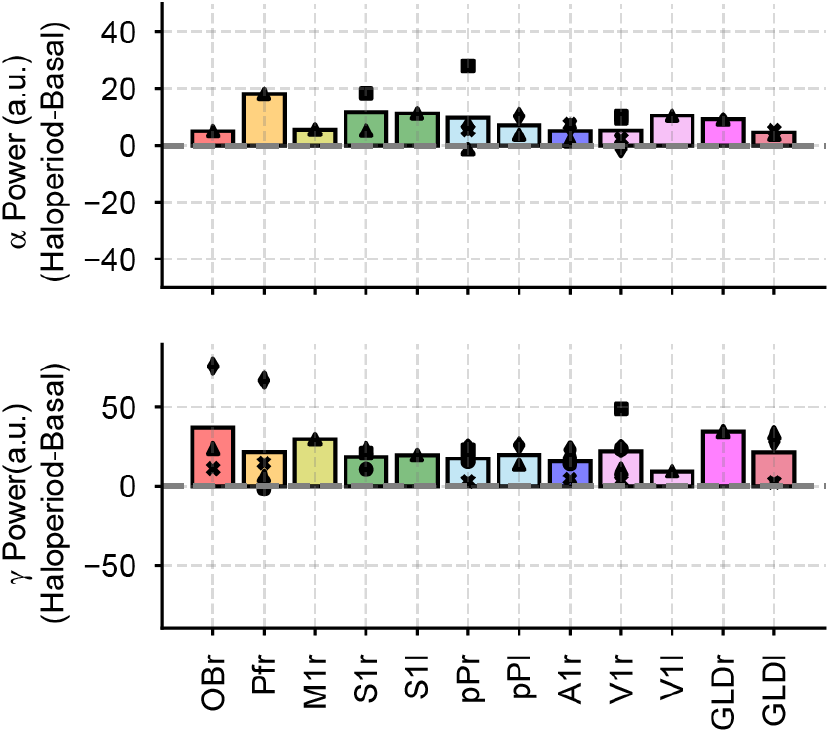
Haloperidol enhances alpha, and gamma total band activity during wakefulness. Bar graph representing the power difference between haloperidol and basal conditions for the whole band for the three frequency bands detected. Each colored bar shows a region in the brain diagrams represented in the figure 3A. Individual points within each bar show an individual animal with a distinct symbol, also shown in figure 3A.

